# Adherence to Life’s Essential 8 Enhances Gut Microbiota Diversity and Cognitive Performance

**DOI:** 10.1101/2025.03.02.641089

**Authors:** Yannick N. Wadop, Jazmyn Muhammad, Rebecca Bernal, Claudia L. Satizabal, Alexa Beiser, Ramachandran S. Vasan, Ramnik Xavier, Tiffany Kautz, Sudha Seshadri, Jayandra Jung Himali, Bernard Fongang

## Abstract

Emerging evidence suggests a complex interplay among cardiovascular health, gut microbiome composition, and cognitive function. Life’s Essential 8 (LE8), developed by the American Heart Association, includes vital metrics of cardiovascular health, such as diet, physical activity, nicotine exposure, sleep health, body mass index (BMI), blood glucose, blood lipids, and blood pressure. In this study, we analyzed data from 781 participants in the Framingham Heart Study (FHS) to explore the relationship between LE8 adherence, gut microbiota, and cognitive performance. Participants with greater adherence to LE8 demonstrated significantly increased gut microbial diversity (α-diversity: Chao1, p = 0.0014; Shannon, p = 0.0071) and distinct microbial compositions (β-diversity: PERMANOVA p = 1e-4). Higher adherence to LE8 was related to an increased abundance of genera *Barnesiella* and *Ruminococcus*, while a reduced abundance of *Clostridium* was associated with higher LE8 adherence. Greater gut microbial diversity (α-diversity: Chao1, p = 0.0012; Shannon, p = 0.0066), and beneficial genera like *Oscillospira* correlated with better global cognitive scores (GCS). Taxonomic overlap analyses revealed microbial taxa that simultaneously influence both LE8 adherence and cognitive outcomes. Mediation analyses indicated that specific taxa, including *Barnesiella* and *Lentisphaerae*, mediated the link between LE8 adherence and cognitive performance. These taxa may serve as key modulators in the gut-brain axis, connecting cardiovascular and brain health. Conversely, higher Clostridium abundance was associated with poorer cognitive performance. This study highlights the significance of comprehensive cardiovascular health metrics in shaping gut microbiota and enhancing cognitive resilience. Our findings underscore the therapeutic potential of targeting gut microbiota to mitigate cognitive decline, warranting further exploration through longitudinal and metagenomic studies.

## Introduction

The interconnectedness of cardiovascular health, gut microbiota, and cognition has emerged as a critical focus of biomedical research. Life’s Essential 8 (LE8), established by the American Heart Association, provides a comprehensive framework to optimize cardiovascular health through key metrics: diet, physical activity, nicotine exposure, sleep health, BMI, blood glucose, blood lipids, and blood pressure.^1^ Adherence to LE8 has been associated with reduced risks of cardiovascular disease, stroke, and neurodegenerative conditions, making it an essential metric for overall health.^2–5^

Recent studies have highlighted significant links between cardiovascular health and cognitive outcomes.^6–8^ Poor cardiovascular health, characterized by risk factors such as hypertension, obesity, and poor sleep, has been implicated in cognitive decline and dementia.^9–12^ Optimal adherence to LE8 metrics has been correlated with improved cognitive performance, likely due to enhanced blood flow, reduced neuroinflammation, and mitigation of vascular risk factors.^3,13,14^ For instance, higher adherence to LE8 has been linked to better executive function and memory performance, underscoring the importance of cardiovascular health in maintaining cognitive resilience.^3,15^

The gut microbiome—a complex ecosystem of trillions of microorganisms^16,17^—plays a vital role in human health, influencing metabolism, immunity, and neural processes.^18–21^ Individual components of LE8, such as diet and physical activity, are known to profoundly shape the gut microbiota.^22^ Diets rich in fiber and polyphenols promote the growth of beneficial taxa like *Faecalibacterium* and *Bifidobacterium*,^23–26^ while physical activity has been shown to enhance microbial diversity and abundance of short-chain fatty acid (SCFA)-producing bacteria.^27–30^ Conversely, suboptimal adherence to LE8, such as diets high in saturated fats and sedentary lifestyles, promotes dysbiosis, characterized by reduced microbial diversity and an overrepresentation of pathogenic taxa.^31–33^ This dysbiosis is increasingly recognized as a precursor to systemic inflammation and chronic disease.^34,35^

The gut-brain axis provides a mechanistic link between the gut microbiome and cognitive function.^36^ Microbial metabolites, such as SCFAs, play neuroprotective roles by reducing inflammation, enhancing the integrity of the blood-brain barrier, and modulating neurotransmitter synthesis.^37–39^ Dysbiosis has been implicated in cognitive impairment through mechanisms involving systemic inflammation, oxidative stress, and altered neurochemical signaling.^40,41^ Emerging research suggests that the gut microbiome may mediate the association between LE8 adherence and cognitive outcomes. For example, individuals adhering to healthy lifestyle practices (measured by LE8) exhibit microbiomes enriched with taxa associated with neuroprotection, such as *Barnesiella* and *Ruminococcus*.^22^ These taxa are thought to modulate brain function via SCFA production and anti-inflammatory pathways.

Despite these associations, the interplay between LE8, the gut microbiome, and cognition remains underexplored. Understanding whether and how the gut microbiome mediates the LE8-cognition link could reveal novel therapeutic targets and inform strategies for optimizing both cardiovascular and brain health. This study aims to elucidate these relationships, hypothesizing that higher adherence to LE8 is associated with a more diverse and balanced gut microbiome, which mediates its protective effects on cognition.

## Methods

### Study Design and Participants

This cross-sectional study utilized data from the Framingham Heart Study (FHS), a longitudinal cohort initiated in 1948 to investigate cardiovascular disease and its risk factors.^42,43^ The FHS has expanded to include multiple generations, providing comprehensive health data over decades. This study included FHS participants from Generation 3, New Offspring Spouses, and Omni2, who provided stool samples and participated in clinical interviews, laboratory tests, health questionnaires, and physical examinations at the third examination (2016-2019). Participants who didn’t provide the required information necessary to compute life’s essential adherence scores were excluded.

### Assessment of Life’s Essential 8 (LE8) Metrics

The American Heart Association (AHA) introduced Life’s Essential 8 (LE8) in 2022, which encompasses eight cardiovascular health metrics: diet, physical activity, nicotine exposure, sleep health, body mass index (BMI), blood glucose, blood lipids, and blood pressure. Each metric is scored on a scale from 0 to 100, with higher scores indicating better health status. The overall LE8 score was derived based on the AHA guidelines^1^, with modifications made to fit the FHS cohorts.^44^ Briefly, diet score was calculated using Dietary Approaches to Stop Hypertension (DASH) based on components including the intake of vegetables, fruits, nuts, legumes, and whole grains; as well as low-fat dairy, and the intake of red and processed meat, sugar-sweetened beverages, and sodium.^45^ The physical activity score was computed by considering the duration and strength of activities including sleep, sedentary, and different strengths of activities (i.e. light, moderate, and vigorous).^46^ The other scores such as sleep quality, blood sugar levels, BMI, blood lipid, blood pressure were derived based on the standard of AHA.^1^ The overall LE8 score was calculated as the unweighted average of the eight metrics.

Based on AHA-established cutoffs, participants were categorized into three adherence levels: low (scores 0–49), moderate (scores 50–79), and high (scores 80–100). These thresholds align with AHA guidelines for cardiovascular health assessment. In the study population only 13 individuals had scores < 50. Because of few individuals in low LE8 adherence and for statistical reasons, we combined moderate and low groups into a unique group noted ModLow. So, we finally classified participants into two categories: high (LE8 scores between 80 and 100) and ModLow (LE8 scores below 80).

### Gut Microbiome Collection and Analysis

Stool samples were collected from FHS participants following standardized procedures to ensure sample integrity as previously described.^42,47,48^ Samples were immediately frozen and stored at −80°C until analysis. Microbial DNA was extracted using the Qiagen PowerSoil DNA Isolation Kit. The V4 region of the 16S rRNA gene was amplified and sequenced on the Illumina MiSeq platform, generating paired-end reads. Sequence data were processed using the DADA2 pipeline^47^ to infer exact sequence variants, followed by taxonomic assignment against the SILVA database. Alpha diversity metrics (Chao1, Shannon, Simpson) were calculated to assess microbial richness and evenness, while beta diversity was evaluated using Bray-Curtis dissimilarity and visualized through principal coordinates analysis (PCoA).

### Global Cognitive Score Assessment

To assess cognitive performance, participants completed cognitive assessments that were administered by trained neuropsychological raters using a comprehensive and standardized test battery during FHS examination cycles as previously described.^49^ The current study included the tests such as trail making test part B (measuring processing speed and executive functions), logical memory (measuring immediate and delayed recall; verbal memory), and visual reproduction (measuring immediate and delayed recall; visual memory) from Wechsler Memory Scale,^50^ and the similarities (measuring executive functions) from the Wechsler Adult Intelligence Scale.^51^ Trail making test part B scores were naturally log-transformed to normalize its distribution and the logged Trail B scores were multiplied by −1 so that higher scores indicated better performances across all cognitive tests. We assessed global cognition through a global cognitive score derived from principal component analysis generating a single component PC1, as previously indicated.^52–55^

### Study context, hypotheses, and objectives

Figure 1. illustrates our study goal to elucidate the mechanisms underlying the connection between LE8 adherence and cognitive performance through the gut microbiome. Our proposed research tackles this interest in three objectives: (1) assess the association between LE8 adherence and cognitive performance; LE8 adherence and each LE8 component may show association with GCS. (2) Estimate the relationship between gut microbiome and LE8 adherence as well as LE8 components; LE8 adherence may impact the gut microbiome and change its composition. (3) Examine the relationship between gut microbiome and GCS; the gut microbiome may interact bidirectionally with cognition. We hypothesize that the gut microbiome may mediate the association between LE8 and cognitive performance through specific bacteria, emphasizing the mediating role of gut microbiota in the LE8-cognition relationship.

**Figure 1.**
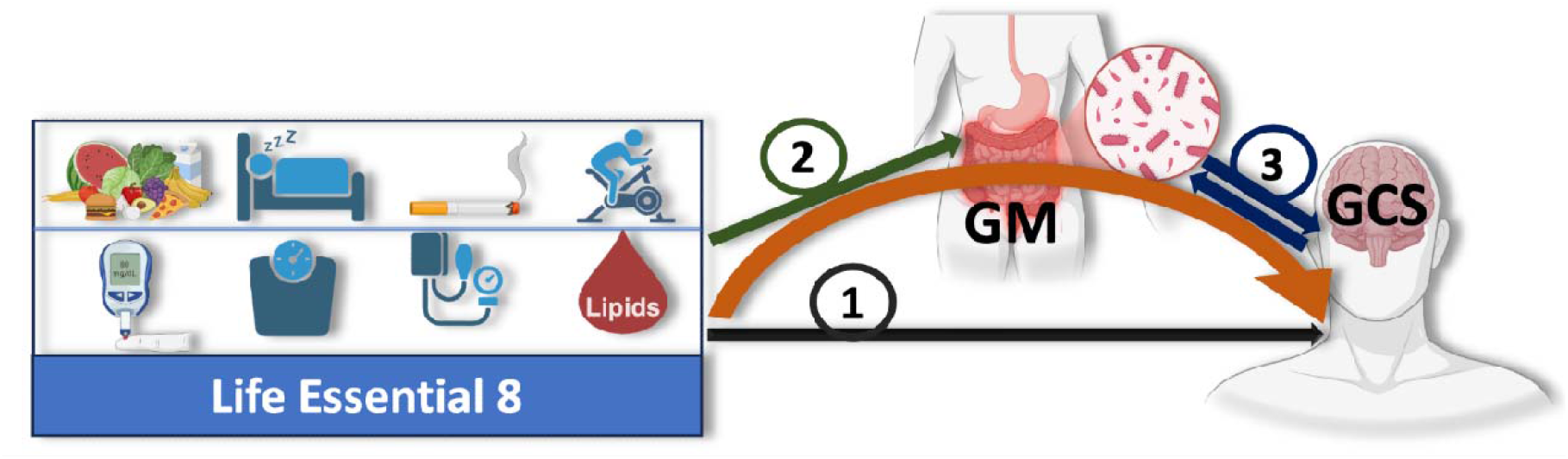
Study Design. Illustration of the hypothesized mediating role of the gut microbiome (GM) in the relationship between adherence to Life’s Essential 8 (LE8) and cognition, measured by the Global Cognitive Score (GCS). The study is structured around three key objectives: (1) assess the relationship between overall LE8 adherence, its individual components, and GCS; (2) evaluate how LE8 adherence and its components influence gut microbiome composition and diversity; and (3) explore bidirectional interactions between the gut microbiome and cognition. The gut microbiome is proposed to mediate the association between LE8 adherence and GCS, with specific microbial taxa or pathways potentially driving this relationship. The orange arrow highlights the central mediating role of the gut microbiota in the LE8-cognition link.

### Statistical Analysis

#### Multivariable association analysis

To evaluate associations between LE8 adherence, gut microbiome diversity, and cognitive performance, we fit general linear models which were implemented using MaAslin2 R package. We also set the MaAlin2 function parameters such as the minimum abundance cutoff at 10^−3^, the minimum prevalence i.e. the minimum percent of samples for which a feature is detected at minimum abundance was at 10%, the negative binomial as the analysis method, and we selected Benjamini-Hochberg to correct the associations’ p-value for multiple testing. The association between cognitive performance and gut microbiome was conducted using the following model,

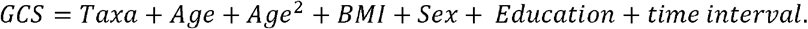

Taxa represents the relative abundance of counts and time interval represents the time difference between stool collection and cognitive tests. However, the following model was used to conduct the association between LE8 adherence and gut microbiome,

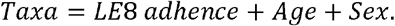

The relationship between LE8 adherence and cognitive performance was carried out using generalized linear model implemented in R software after controlling for age and sex. Statistical significance was set at p < 0.05, with Benjamini-Hochberg correction applied for multiple comparisons.

### Microbiome diversity analysis

We estimated the alpha-diversity using Chao1 and Shannon indexes, which are used to assess difference in species richness and evenness between LE8 adherence groups. Microbiome data were also analyzed using the *vegan*^56^ package in R, with permutational multivariate analysis of variance (PERMANOVA) employed to assess differences in beta diversity.

### Mediation analysis

To explore whether the gut microbiome through specific bacterial taxa mediate the association between LE8 adherence and cognitive performance, we performed a mediation analysis using the R package mediation^57^. The analysis was proceeded in two steps. We used two statistical models; first, the mediator model (*Taxa* = *LE*8 *adherence* + *Age* + *sex*) and the outcome model (*GCS* = *Taxa* + *LE*8 *adherence* + *Age* + *Sex* + *Education*). Both models were fitted separately using generalized linear model implemented in R software. Secondly, the fitted objects including the main inputs were used as arguments in mediation function in R to compute the average causal mediation effects (indirect effects) and average direct effects (direct effects). Bootstrapping was implemented in mediation function to estimate if the mediation effects are statistically significant.

## Results

### Participant Characteristics

The study cohort comprised participants aged 32 to 78 years, with a mean age of 54.9 years. The majority of participants were female (57.1%), and 64.4% had completed college-level education. The median Life’s Essential 8 (LE8) score was 76.9. Participants classified in the high adherence group demonstrated significantly superior cognitive performance compared to those in the combined moderate and low adherence group (p < 0.001). Demographic distributions showed no significant differences in gender or baseline health status between groups, underscoring the robust association of LE8 adherence with outcomes.

**Table 1.** Demographic Characteristics.

| Variables | Overall (N = 781) | High LE8 (N = 322) | ModLow (N = 459) | P-value |
| --- | --- | --- | --- | --- |
| Age, years | 54.9 ± 8.3 | 53.2 ± 8.5 | 56.2 ± 7.9 | 1.00e-6 |
| Female, n (%) | 446 (57.1) | 227 (70.5) | 219 (47.7) | 0.64 |
| Education, n (%) |  |  |  |  |
| High school degree | 76 (9.7) | 15 (4.7) | 61 (13.3) | 2.88e-13 |
| Some college | 202 (25.9) | 57 (17.7) | 145 (31.6) | 2.20e-16 |
| College graduated | 503 (64.4) | 250 (77.6) | 253 (55.1) | 0.89 |
| Time interval between stool collection and cognitive test, weeks | 133.9 ± 124.2 | 131.2 ± 120.5 | 135.9 ± 126.8 | 0.59 |
| **Cardiovascular risk factors** |  |  |  |  |
| Triglycerides, mg/dL | 110.9 ± 78.3 | 77.6 ± 32.2 | 132.9 ± 92.5 | 2.00e-16 |
| Systolic blood pressure, mmHg | 119 ± 14 | 113 ± 12 | 123 ± 14 | 2.00e-16 |
| Diastolic blood pressure, mmHg | 75 ± 8 | 72 ± 7 | 78 ± 9 | 2.00e-16 |
| Total cholesterol, mg/dL | 190.1 ± 35.0 | 185.1 ± 31.1 | 193.6 ± 37.1 | 5.16e-4 |
| HDL cholesterol, mg/dL | 61.9 ± 19.8 | 69.5 ± 19.2 | 56.5 ± 18.4 | 2.00e-16 |
| Treatment for hypertension, n (%) | 179 (22.9) | 24 (13.4) | 155 (86.6) | 2.20e-16 |
| Stage I hypertension, n (%) | 221 (28.3) | 32 (14.5) | 189 (85.5) | 2.20e-16 |
| Prevalent of cardiovascular disease, n (%) | 30 (3.8) | 10 (33.3) | 20 (66.7) | 2.01e-2 |
| Current smoking, n (%) | 32 (4.1) | 1 (3.1) | 31 (96.9) | 4.17e-13 |
| History of Diabetes, n (%) | 58 (7.4) | 1 (1.7) | 57 (98.3) | 2.20e-16 |
| Body mass index, kg/m<sup>2</sup>, median [Q1, Q3] | 26.8 [24.1, 30.6] | 24.2 [22.1, 26.3] | 29.5 [26.4, 32.8] | 2.20e-16 |
| Global cognition | 0.59 [0.12, 1.09] | 0.71 [0.28, 1.18] | 0.47 [0.01, 0.45] | 2.16e-6 |
| Trail Making part B | 0.07 [-0.17, 0.31] | 0.11 [-0.09, 0.35] | 0.03 [-0.19, 0.24] | 5.00e-5 |
| Similarities | 17.88 ± 2.89 | 18.05 ± 2.79 | 17.77 ± 2.97 | 0.19 |
| Visual Reproduction | 19.04 ± 4.72 | 19.89 ± 4.38 | 18.44 ± 4.85 | 1.60e-5 |
| Logical Memory | 24.79 ± 6.66 | 25.71 ± 6.92 | 24.15 ± 6.41 | 1.50e-3 |
| Life's essential eight scores | 76.3 ± 11.9 | 87.5 ± 5.2 | 68.4 ± 8.4 | 2.00e-16 |
| Diet scores | 45.2 ± 30.2 | 62.3 ± 27.7 | 33.2 ± 25.8 | 2.00e-16 |
| Sleep health scores | 89.9 ± 18.2 | 94.7 ± 12.2 | 86.6 ± 20.8 | 1.70e-11 |
| Nicotine exposure scores | 87.4 ± 22.3 | 94.3 ± 11.9 | 82.5 ± 26.3 | 2.00e-16 |
| Physical activity scores | 83.2 ± 27.9 | 90.9 ± 16.9 | 77.8 ± 32.6 | 6.40e-13 |
| Blood pressure scores | 72.3 ± 28.4 | 87.3 ± 20.4 | 61.8 ± 28.5 | 2.00e-16 |
| Blood glucose scores | 92.3 ± 19.4 | 98.3 ± 8.3 | 88.1 ± 23.4 | 2.00e-16 |
| Blood lipids scores | 72.9 ± 26.9 | 85.5 ± 21.1 | 64.0 ± 27.2 | 2.00e-16 |
| Body mass index scores | 66.9 ± 30.6 | 86.8 ± 18.5 | 53.0 ± 29.8 | 2.00e-16 |

Values are mean±SD, or median [quartile 1, quartile 3] for nonnormally distributed variables. ModLow indicates LE8 group combining individuals with low and moderate LE8 adherence. The tests of difference between both LE8 groups were based on t-test for continuous variables or test of proportion for categorical variables.

### Adherence to LE8 is Associated with Increased Gut Microbial Diversity

Alpha diversity indices were significantly higher in the high LE8 adherence group (Chao1: p = 0.0014; Shannon: p = 0.0071), indicating richer and more even microbial communities (**Figure 2a**). Beta diversity analysis showed distinct clustering of microbial profiles between high LE8 adherence group and the one combining moderate and low, ModLow group (PERMANOVA p = 1e-4), reflecting differences in overall microbial composition (**Figure 2b**). Key phyla observed included *Firmicutes, Bacteroidetes*, and *Proteobacteria*, with the high LE8 adherence group showing a relative increase in beneficial taxa such as *Faecalibacterium* and *Ruminococcaceae* (**Figure S1**).

**Figure 2.**
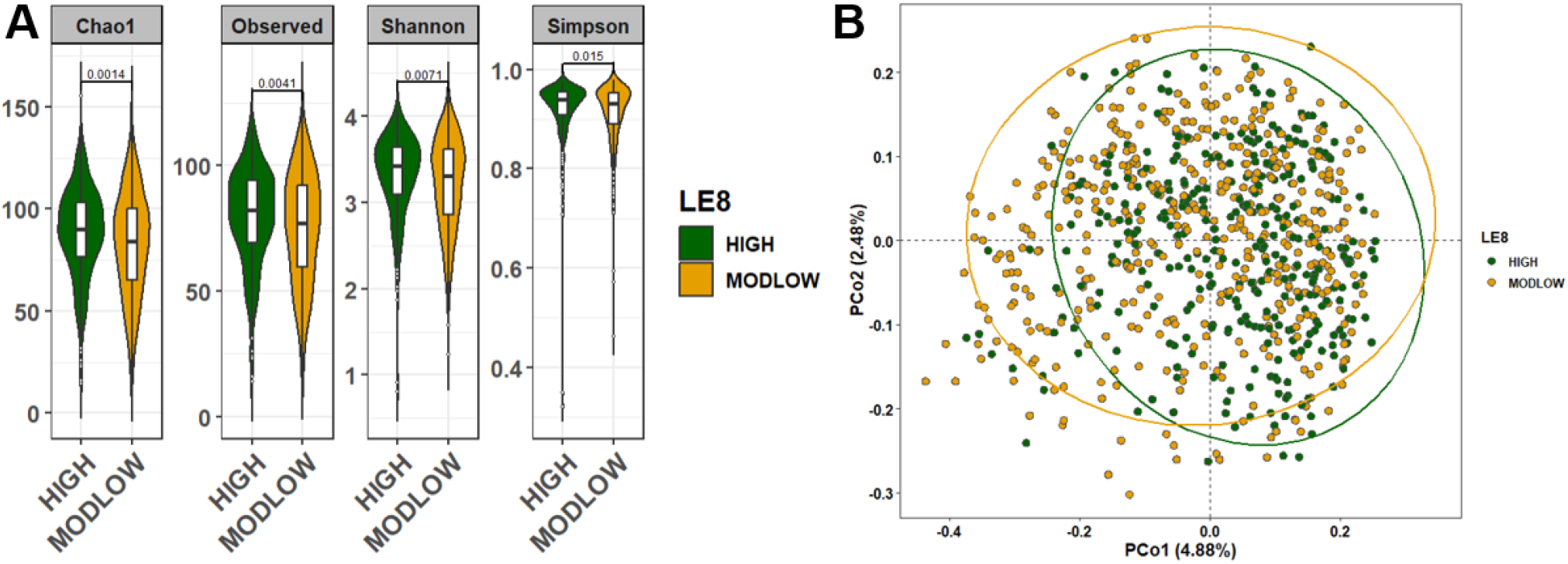
Comparison of the microbial community diversity of samples between high and below high (ModLow) LE8 adherence groups. (A) The α-diversity analysis through calculation of Chao1, Observe, Shannon, and Simpson indexes. The tests of difference in microbial diversity at the OTU level between both LE8 groups were based on Wilcoxon test. (B) Beta diversity on samples represented by PCoA using Bray-Curtis dissimilarity. The p-value of PERMANOVA test is 1.0e-4.

For sensitivity analysis, we excluded individuals with low LE8 adherence (LE8 scores < 50). The results showed that alpha diversity indices were always significantly higher in the high LE8 adherence group compared to moderate adherence group (Chao1: p = 0.0078; Shannon: p = 0.024) as illustrated in **Figure S2**. Beta diversity analysis showed that microbial composition between high and moderate LE8 adherence groups was significantly different (PERMANOVA p = 1e-4).

### Adherence to LE8 is Associated with Specific Patterns of Gut Microbial Abundance

To examine the association between the gut microbiome relative abundance and LE8 adherence, we conducted multivariable linear regression analysis using negative binomial regression through MaAslin2 R package after adjusting for age and sex. Considering specific features with BH adjusted p-value < 0.05, we found several associations at the phylum, family, and genus levels (**Figure 3** and **Figure S3**). Findings revealed that LE8 scores exhibited positive associations with the relative abundance of genera such as *Barnesiellaceae Barnesiella* (β[95% CI], p: 0.0263[0.0261,0.0266], 1.4e-297), *Lachnospiraceae Ruminococcus* (0.0128[0.0122, 0.0134], 6.8e-294), *Ruminococcaceae Clostridium* (0.0583[0.0401, 0.0766], 2.9e-9), *Clostridiaceae Clostridium* (0.0283[0.0092, 0.0475], 1.3e-2), *Ruminococcaceae Ruminococcus* (0.0091[0.0018, 0.0163], 4.1e-2), and *Victivallaceae Victivallis* (0.043[0.0153, 0.0707], 9.5e-3). Additionally, LE8 scores showed a negative association with the relative abundance of genera *Lachnospiraceae Clostridium* (−0.0521[−0.0673, −0.0368], 1.9e-10), *Ruminococcaceae Subdoligranulum* (−0.0248[−0.0378, −0.0119], 9.9e-4), *Coriobacteriaceae Collinsella* (−0.0216[−0.0334, −0.0009], 1.8e-3), *Erysipelotrichaceae Clostridium* (−0.0261[−0.0424, −0.0097], 7.7e-3), *Streptococcaceae Streptococcus* (−0.0301[−0.0499, −0.0102], 1.1e-2), *Coriobacteriaceae Eggerthella* (−0.0345[−0.0601, −0.0089], 2.6e-2), and *Erysipelotrichaceae Catenibacterium* (−0.0129[−0.0136, −0.0124], 7.2e-282). These genera associated with LE8 scores belong to some specific families and phyla that showed linkages with LE8 groups (High, ModLow) as well. Notably, the increased abundance of bacteria such as *Methanobacteriaceae, Victivallaceae, Barnesiellaceae*, and *Clostridiaceae* displayed an association with higher LE8 scores. While the decreased abundance of *Streptococcaceae* and *Coriobacteriaceae* had an association with greater LE8 scores at the family level. Moreover, the LE8 scores demonstrated positive relationships with the relative abundance of several phyla, including *Firmicutes, Actinobacteria, Proteobacteria*, and *Lentisphaerae*.

**Figure 3.**
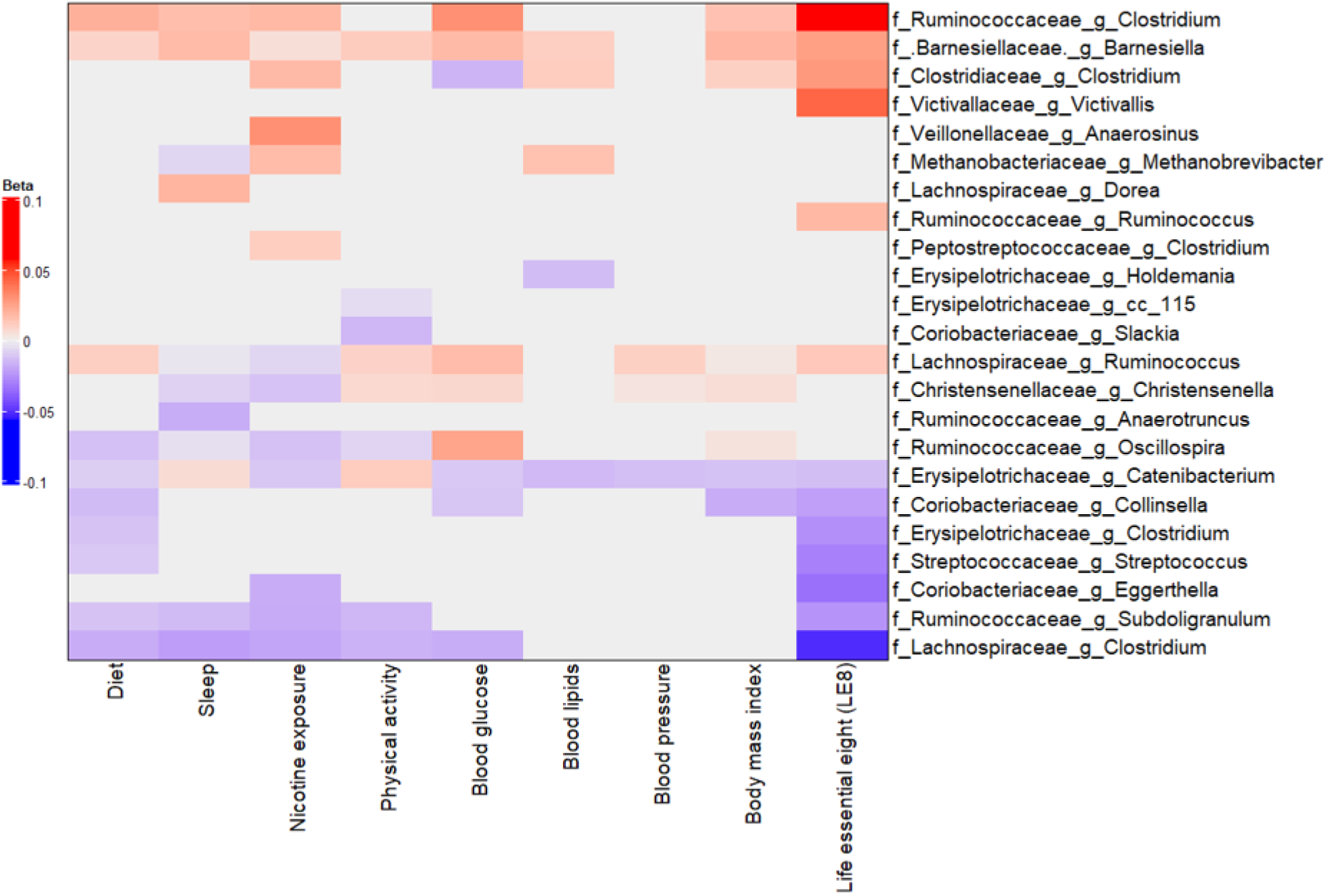
Multivariable association between gut microbiota and LE8 adherence. Heatmap depicting the significant (adjusted p-value < 0.05) genera associated with LE8 adherence along with its components.

**Figure 4.**
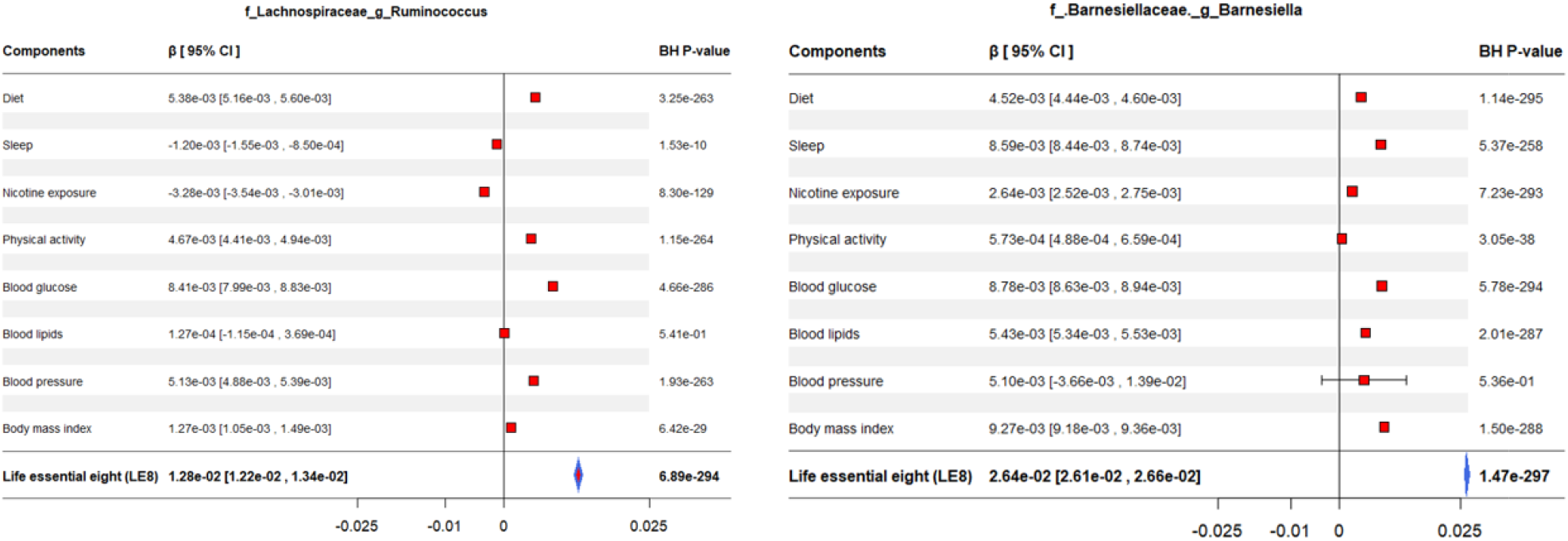
Forest plot displaying the effects of LE8 components in the association between gut microbiome and LE8 adherence. The significant contribution of each metric in the association between gut bacteria and LE8 adherence is clearly observed.

### The Gut Microbiome Associates with Components of LE8

In a secondary analysis, we estimated which components of LE8 may drive the relationship between gut microbiome and LE8 (**Figures 3** and **4**). The results suggested that the top 4 components linked with the gut microbiome were nicotine exposure, BMI, diet, and sleep health. Although few relations were observed with some components, we found that the genus *Barnesiellaceae Barnesiella* showed positive linkages with the components of LE8 except for blood pressure. Its higher abundance was especially highly associated with better scores for these components. The reduced abundance of the bacterium *Lachnospiraceae Ruminococcus* was also highly associated with greater scores of sleep health and nicotine exposure. At the same time, its increase was significantly associated with higher scores for diet, BMI, physical activity, blood glucose, and blood pressure. However, the higher scores for diet, sleep health, nicotine exposure, physical activity, and blood pressure were associated with a reduced abundance of *Lachnospiraceae Clostridium*. Finally, *Ruminococcaceae Clostridium* showed positive relationships with diet, sleep health, nicotine exposure, blood glucose, and BMI. Further links are displayed in **Figure 3** and **S4**.

### Association between the Gut Microbiome Genera and Cognitive Performance

The GCS distribution was equally divided into quintiles. Individuals with GCS within the first quintile (lowest scores) were classified as **poor**, and the remainder (highest scores) were considered **normal**. Participants with higher GCS exhibited increased alpha diversity (Chao1: p = 0.0012; Shannon: p = 0.0066) and distinct microbial profiles (**Figures S5** and **S6**). Beta diversity analysis revealed that microbial composition between normal and poor groups was significantly different (PERMANOVA p = 1e-4).

To estimate the relationship between the relative abundance of gut microflora and GCS, we carried out multivariable linear regression analysis using negative binomial regression after adjusting for age, sex, body mass index, education, and the time interval between stool sample and cognitive tests (**Figure 5** and **Figure S7**). At the genus level, poorer GCS had a statistically significant association with a lower abundance of genera *Lachnospiraceae Ruminococcus* (β[95% CI], p: 0.2523[0.2423, 0.2623], 5.3e-281), *Barnesiellaceae Barnesiella* (0.1219[0.1182, 0.1255], 4.4e-267), *Ruminococcaceae Oscillospira* (0.0943[0.0663, 0.1223], 3.0e-10), and *Victivallaceae Victivallis* (0.6558[0.1840, 1.1277], 2.9e-2). At the same time, these lower GCS showed an association with elevated abundance of *Erysipelotrichaceae cc_115* (−0.0396[−0.0579, −0.0213], 1.3e-4), *Lachnospiraceae Clostridium* (−0.3984[−0.6608, −0.1361], 1.4e-2*), Ruminococcaceae Subdoligranulum* (−0.2950[−0.5155, −0.0775], 3.7e-2), and *Ruminococcaceae Anaerotruncus* (−0.3881[−0.6889, −0.0871], 4.7e-2). The families *Methanobacteriaceae, Coriobacteriaceae, Christensenellaceae, Rikenellaceae, Ruminococcaceae*, and *Victivallaceae* had positive correlations with GCS. Finally, significant associations were observed between poorer GCS and decreased abundance of bacteria *OD1, Lentisphaerae*, and *Firmicutes* at the phylum level. Taxonomic analysis suggested that beneficial genera contribute to neuroprotective mechanisms, potentially mediated through the production of short-chain fatty acids.

**Figure 5.**
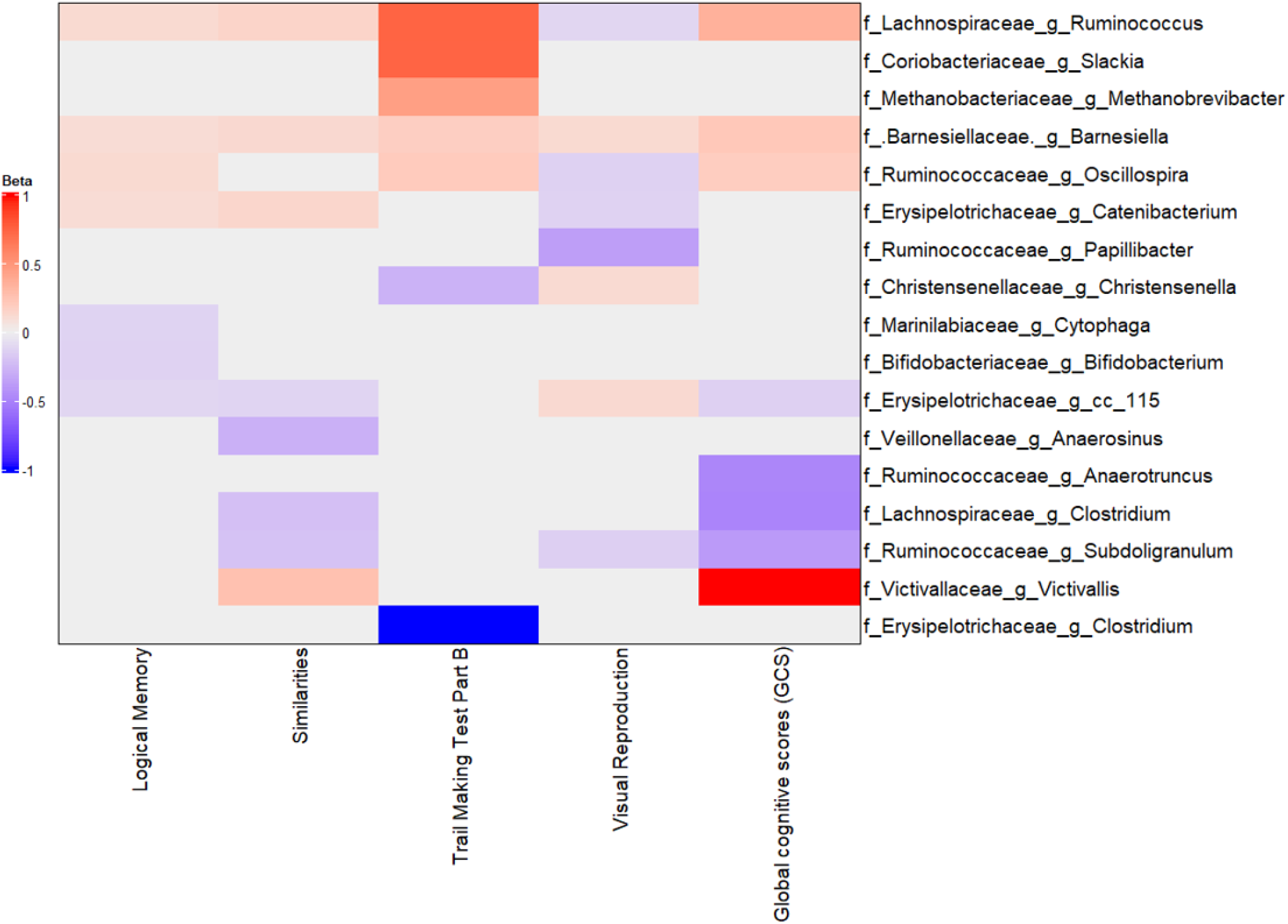
Multivariable association between gut microbiota and global cognitive scores. Heatmap depicting the significant (adjusted p-value < 0.05) genera associated with GCS along with the scores of tests composing GCS.

### The Gut Microbiome Associates with Neuropsychological Tests Composing the GCS

Considering the four tests composing GCS, results suggested that the reduced proportion of bacterium *Barnesiellaceae Barnesiella* that was highly linked with poorer GCS, exhibited a significant association with lower scores of similarities, visual reproduction, logical memory, and trail-making test parts B (**Figure 5**). Findings also suggested that the lower abundance of genus *Lachnospiraceae Ruminococcus* showed an association with higher scores of visual reproduction and lower scores of three other tests. While the weaker abundance of *Lachnospiraceae Clostridium* and *Ruminococcaceae Subdoligranulum* had a link with smaller scores on the similarities test.

### Overlapping taxa among bacteria associated with LE8 and GCS

Having identified the bacteria in relationship with LE8 and GCS, we noticed that there were twelve overlapped taxa among both microbiome profiles. The overlapping taxa included *Barnesiellaceae Barnesiella, Lachnospiraceae Ruminococcus, Lachnospiraceae Clostridium, Ruminococcaceae Subdoligranulum, Victivallaceae Victivallis, Victivallaceae, Methanobacteriaceae, Coriobacteriaceae, Lentisphaerae*, and *Firmicutes*. Suggesting that these bacteria might play dual roles in modulating cardiovascular and cognitive health.

### Adherence to LE8 Significantly Associates with an Increase in Cognitive Performance

Research has previously demonstrated an association between LE8 and cognition.^2,58,59^ Our objective was to assess the association between LE8 adherence and global cognition in the study population. Performing a multivariable linear regression to test the link between GCS and LE8 after adjusting for age and sex, we found that greater LE8 scores had a statistically significant association with higher GCS (β=0.0094, 95% CI [0.0051, 0.0138], adjusted p-value=2.1e-5) as illustrated in **Figure 6**. We also carried out multivariable logistic regression analysis to differentiate the GCS of individuals in both LE8 groups. Findings showed that individuals in the high LE8 group had greater GCS (OR=1.225, 95% CI [1.220, 1.3280]; adjusted p-value=1.2e-4).

**Figure 6.**
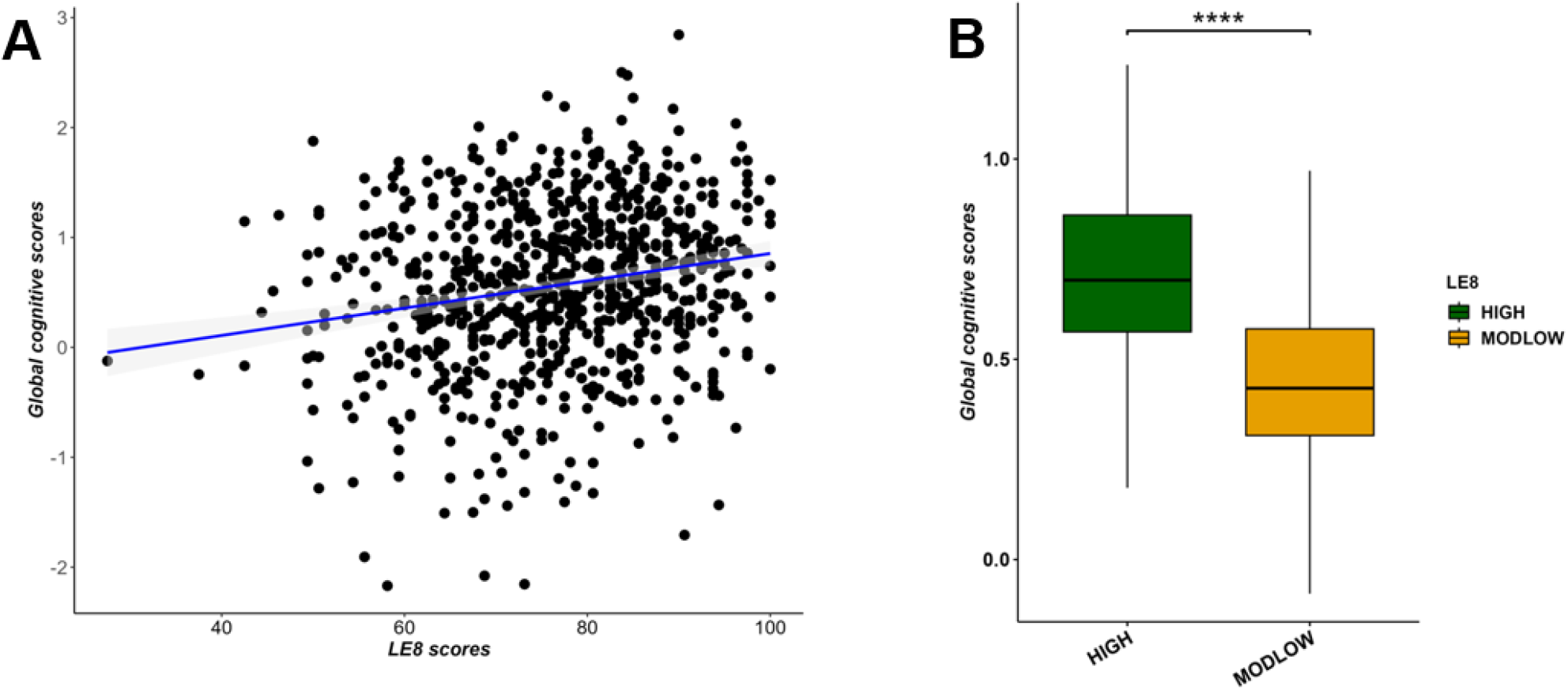
Multivariable association between LE8 adherence and global cognitive scores adjusting for age and sex. (A) scatter plot showing a positive correlation between LE8 adherence and GCS. (B) Boxplot displaying significant differences in global cognitive scores among LE8 groups.

### On the Mediating Role of the Gut Microbiome Between Adherence to LE8 and Cognitive Performance

Focusing on bacteria that overlapped between microbial profiles showing association with LE8 and GCS, we carried out a mediation analysis using a generalized linear model through the *mediation*^57^ R package to identify bacteria that serve as a bridge to the linkage of LE8 with GCS in the FHS population. Mediation analyses revealed that specific bacterial taxa mediated the relationship between LE8 adherence and cognitive performance. Notable mediators included *Victivallaceae Victivallis, Barnesiellaceae Barnesiella, Lachnospiraceae Ruminococcus, Lachnospiraceae Clostridium, Victivallaceae, Methanobacteriaceae, Lentisphaerae*, and *Firmicutes*, which exhibited significant indirect effects, suggesting their role in the gut-brain axis (**Figures 7, 8**, and **S8**). The findings indicated that high LE8 adherence seems to increase the abundance of these bacteria except for *Lachnospiraceae Clostridium*, which consequently increases the scores of composite cognitive tests. Conversely, results revealed that moderate and low LE8 scores seem to increase the level of *Lachnospiraceae Clostridium* with an indirect effect IDE=0.0065, 95% CI [0.004, 0.01], which leads to the reduction of GCS. We then examined the effects with strongest indirect effects (IDE). In **Figure 8**, we observed that the top 4 bacteria with the strongest IDE included *Victivallaceae Victivallis* (IDE=0.0173, 95% CI [0.0143, 0.02]), *Victivallaceae* (IDE=0.0166, 95% CI [0.0139, 0.02]), *Barnesiellaceae Barnesiella* (IDE= 0.0160, 95% CI [0.0130, 0.02]), and *Lentisphaerae* (IDE= 0.0144, 95% CI [0.0118, 0.02]). We also computed the direct effects of these taxa and noticed that there are only few bacteria displaying significant estimated direct effects, as indicated in **Figure S9**. These bacteria included *Firmicutes* (DE=0.0074, 95% CI [0.0028, 0.01]), *Methanobacteriaceae* (DE=0.0060, 95% CI [0.0014, 0.01]), *Lachnospiraceae Clostridium* (DE=0.0059, 95% CI [0.0024, 0.01]), *Victivallaceae* (DE=-0.0041, 95% CI [−0.0075, 0.0]), and *Victivallaceae Victivallis* (DE=-0.0049, 95% CI [−0.0082, 0.0]).

**Figure 7.**
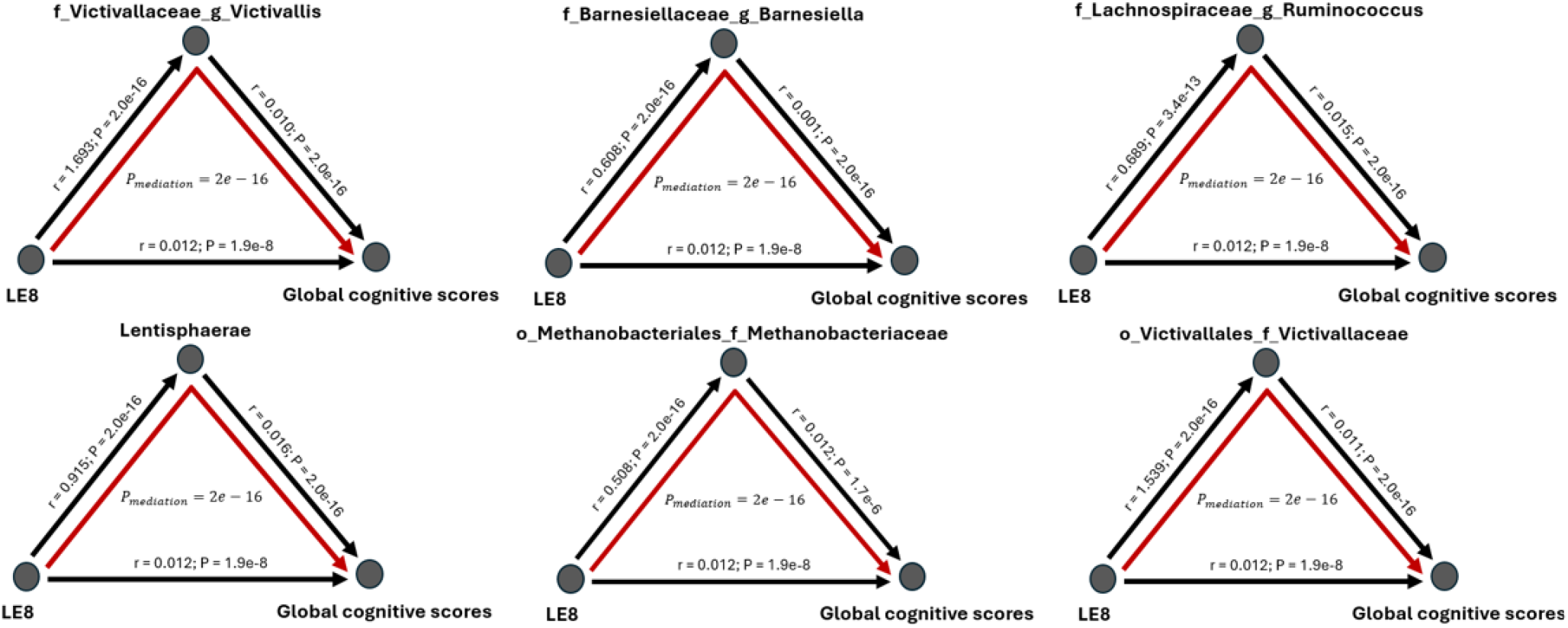
Mediation analysis of gut microbiome on the association between LE8 adherence scores and global cognitive scores. The mediation effect of bacteria *Victivallaceae Victivallis, Barnesiellaceae Barnesiella, Lachnospiraceae Ruminococcus, Lachnospiraceae Clostridium, Victivallaceae, Methanobacteriaceae* on association of lower cognition with moderate and low LE8 adherence.

**Figure 8.**
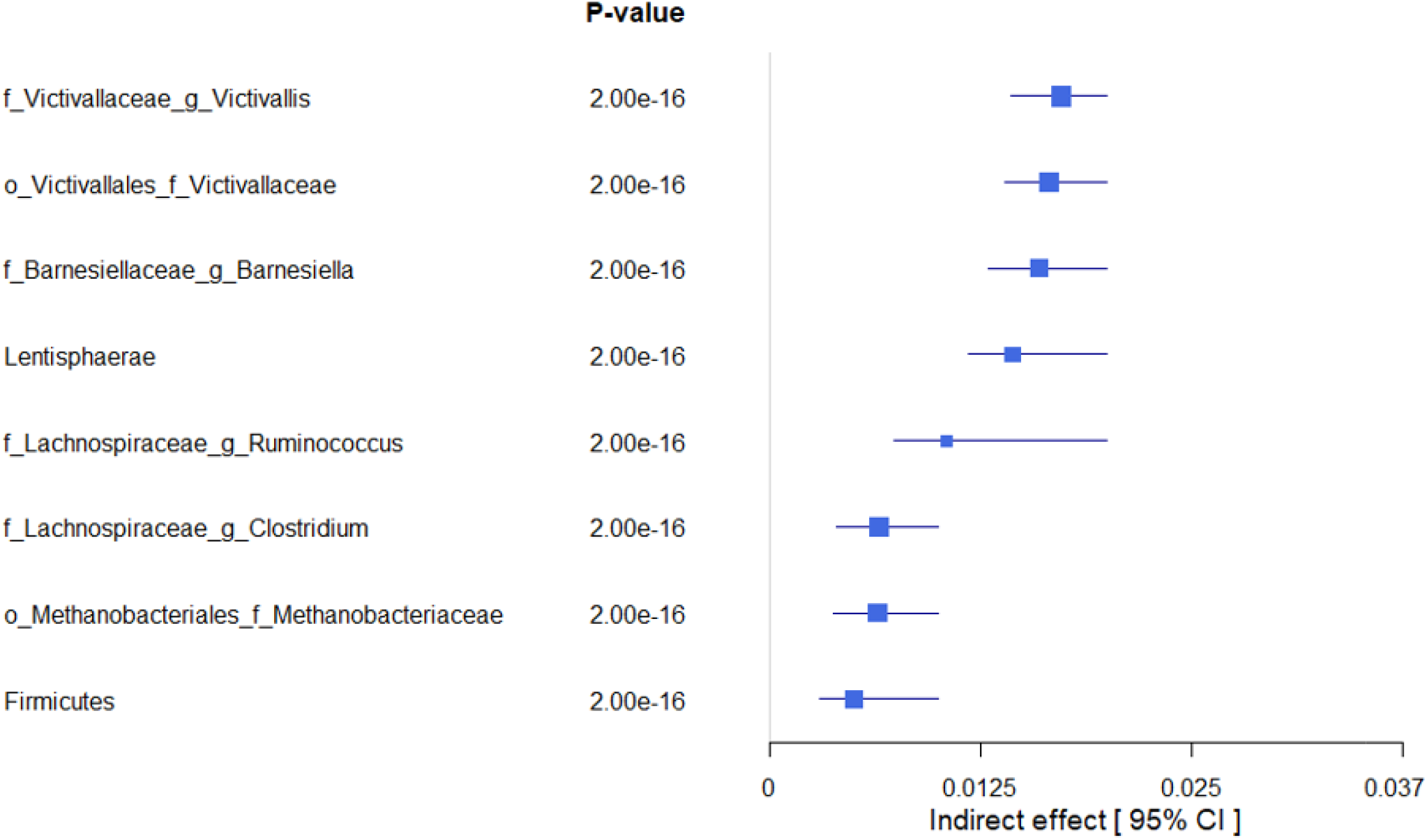
Forest plot showing the estimated indirect effects of bacteria that mediate the association between higher adherence to LE8 with better cognition. As we can see, the top 4 strongest IDE include *Victivallaceae Victivallis, Victivallaceae, Barnesiellaceae Barnesiella, and Lentisphaerae*.

## Discussion

This study provides a comprehensive examination of the relationships between adherence to Life’s Essential 8 (LE8), gut microbiome, and cognitive performance, utilizing robust data from the Framingham Heart Study. Our findings emphasize the critical role of cardiovascular health metrics in shaping gut microbiota and their downstream effects on cognitive resilience.

Our results demonstrate that higher LE8 adherence is associated with greater microbial diversity and evenness, as evidenced by significant differences in alpha diversity indices between high and moderate adherence groups. The increase in beneficial taxa such as *Faecalibacterium* and *Barnesiella* suggests that LE8 adherence promotes a gut microbiota composition favorable for health. Concurrently, beta diversity analysis revealed distinct clustering of microbial profiles, indicating that LE8 adherence influences overall microbial community structure.

The mediation analysis highlights the gut microbiome as a key intermediary linking LE8 adherence to cognitive performance. Specifically, taxa such as *Barnesiella* and *Ruminococcus* were identified as significant mediators. These genera are known for their anti-inflammatory properties and production of short-chain fatty acids, which are critical for maintaining gut-brain axis integrity.

The findings have significant implications for public health and clinical practice. LE8 metrics represent modifiable lifestyle factors that can be targeted through behavioral interventions to improve both cardiovascular and cognitive outcomes. The identification of specific microbial taxa as mediators underscores the potential for microbiome-based diagnostics and therapeutics. For example, probiotics or dietary interventions designed to enrich beneficial taxa could serve as adjunctive strategies for cognitive health maintenance.

We showed the interconnected nature of cardiovascular health, gut microbiota, and cognition. By adhering to LE8 metrics, individuals not only improve cardiovascular health but also potentially enhance cognitive resilience through microbiome-mediated mechanisms. This underscores the importance of a holistic approach to health that integrates cardiovascular and neurological perspectives.

While the study provides valuable insights, it also raises important questions for future research. Longitudinal studies are needed to establish causal relationships among LE8 adherence, microbiome diversity, and cognitive outcomes. Advanced sequencing techniques, such as metagenomics, should be employed to investigate functional pathways within the microbiome. Additionally, randomized controlled trials could validate the efficacy of microbiome-targeted interventions in enhancing cognitive performance.

## Conclusion

This study highlights the interconnected relationship between cardiovascular health, gut microbiome diversity, and cognitive function. Higher adherence to Life’s Essential 8 metrics was associated with favorable microbial profiles and enhanced cognitive performance, with the gut microbiome serving as a critical mediator. These findings emphasize the importance of integrated lifestyle interventions that address cardiovascular and cognitive health simultaneously. To validate these results and refine therapeutic strategies, future research should prioritize longitudinal studies and randomized controlled trials that explore the causal pathways and clinical applications of these findings. By targeting modifiable factors such as diet, physical activity, and microbiome health, this research underscores a proactive approach to optimizing both physical and mental health outcomes.

## Acknowledgements

The Authors thank Dr. Ramnik Xavier from the Broad Institute of MIT and Harvard, and the Center for Microbiome Informatics and Therapeutics (Massachusetts Institute of Technology, Cambridge, MA, USA) for providing access to the FHS microbiome data.

