## Supplementary file for "Adherence to Life’s Essential 8 Enhances Gut Microbiota Diversity and Cognitive Performance"

**Taxonomic profiling from stool samples of FHS subjects in LE8 adherence groups**

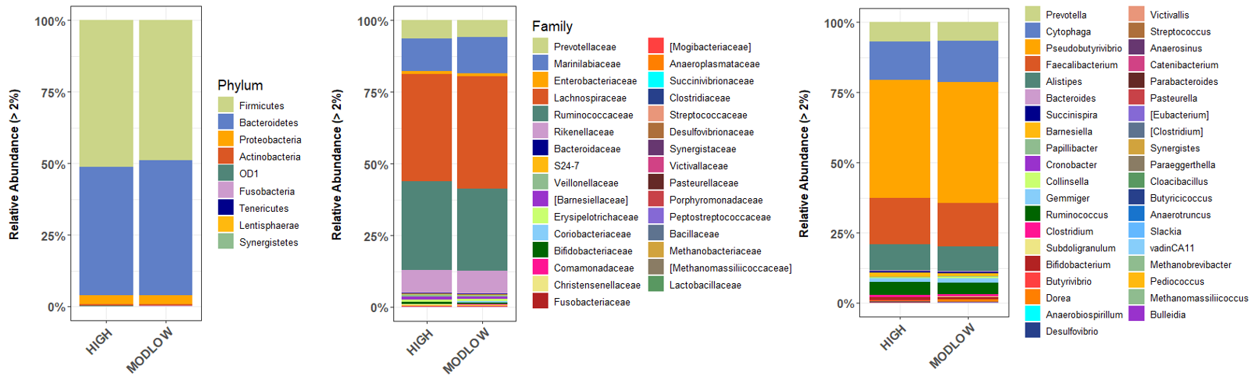

**Figure S1.** Stacked plot showing the microbiome profiles of predominant taxa in the stool samples of high and ModLow LE8 adherence groups at the phylum, family, and genus levels (from left to right).

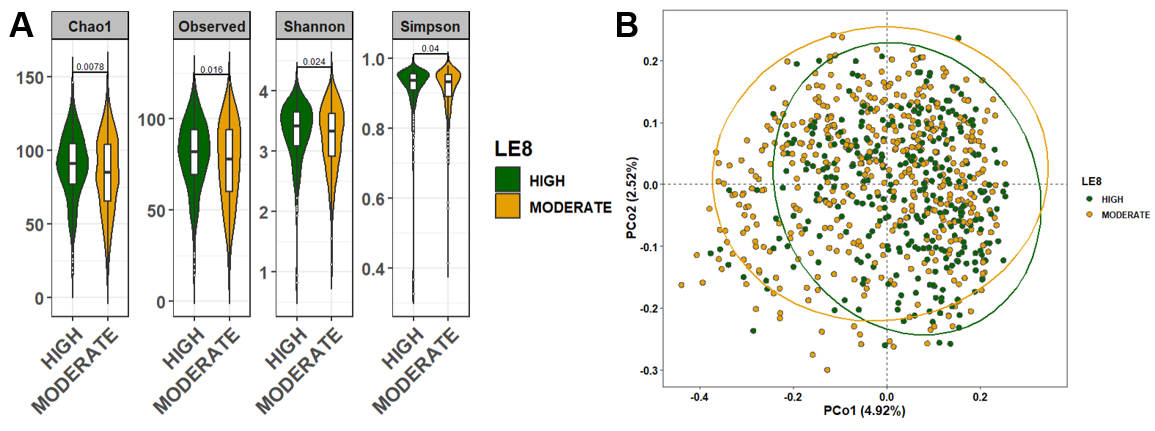

**Figure S2.** Comparison of the microbial community diversity of samples between high and moderate LE8 adherence groups. (A) The α-diversity analysis through calculation of Chao1, Observe, Shannon, and Simpson indexes. The tests of difference in microbial diversity at the OTU level between both LE8 groups were based on Wilcoxon test. (B) Beta diversity on samples represented by PCoA using Bray-Curtis dissimilarity. The p-value of PERMANOVA test is 1.0e-4.

**LE8 Adherence and Gut Microbiome**

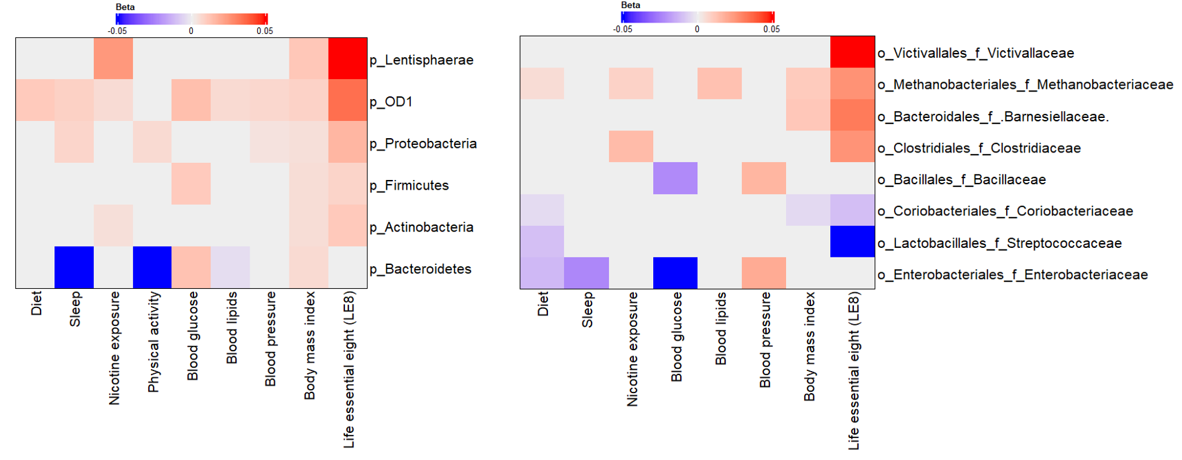

**Figure S3.** Multivariable association between gut microbiota and LE8 adherence. Heatmap depicting the significant (adjusted p-value < 0.05) taxa associated with LE8 adherence along with its components at the phylum (left) and family (right).

**Forest plots exhibiting the effects of LE8 components on LE8-gut microbiome relationships**

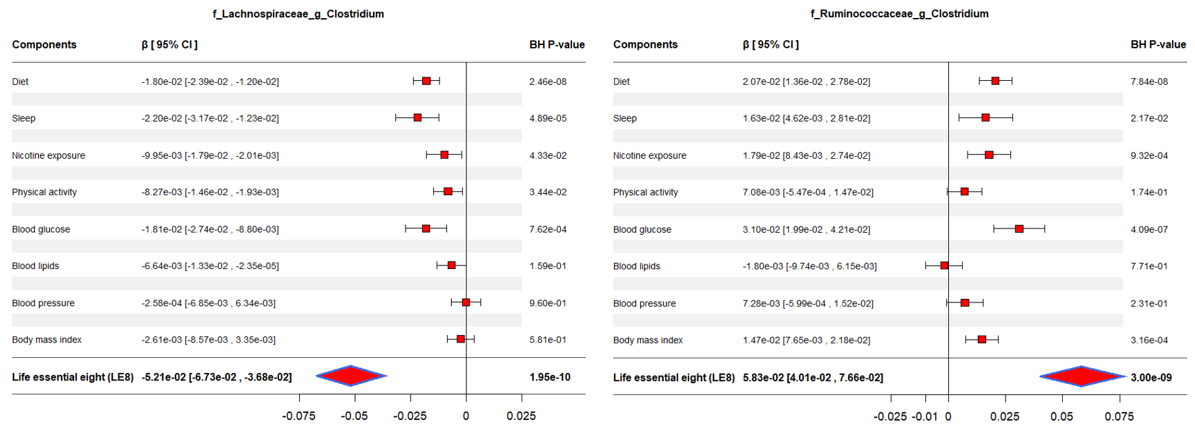

**Figure S4.** Forest plot displaying the effects of LE8 components in the association between gut microbiome and LE8 adherence.

**Diversity analysis showed differences across GCS groups**

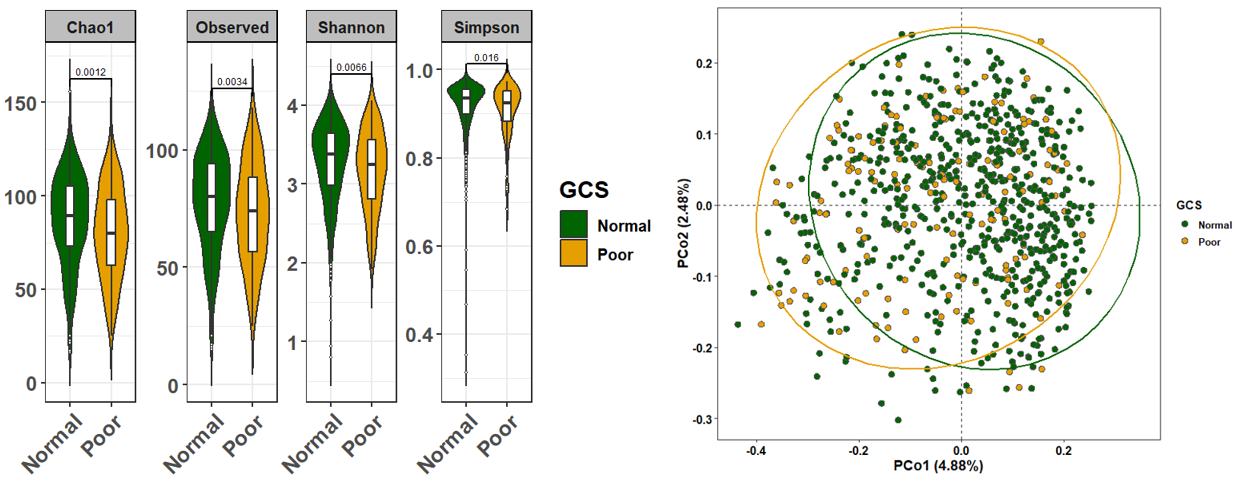

**Figure S5.** Comparison of the microbial community diversity of samples between normal and poor groups of GCS. (A) The α-diversity analysis through calculation of Chao1, Observe, Shannon, and Simpson indexes. The tests of difference in microbial diversity at the OTU level between both GCS groups were based on Wilcoxon test. (B) Beta diversity on samples represented by PCoA using Bray-Curtis dissimilarity. The p-value of PERMANOVA test is 1.0e-4.

**Taxonomic profiling from stool samples of FHS subjects in GCS groups**

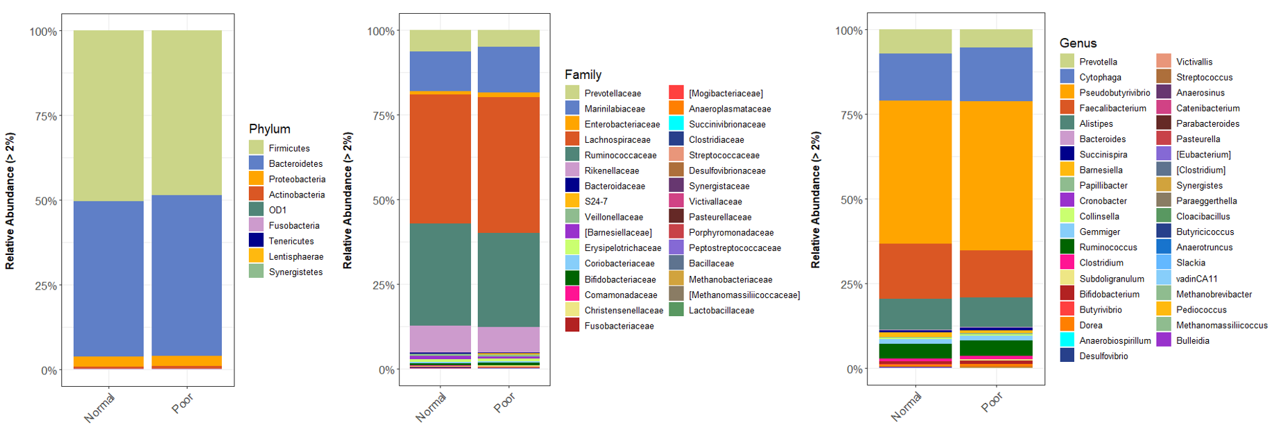

**Figure S6.** Stacked plot showing the microbiome profiles of predominant taxa in the stool samples of normal and poor GCS groups at the phylum, family, and genus levels (from left to right).

**Gut Microbiome and Cognitive Performance**

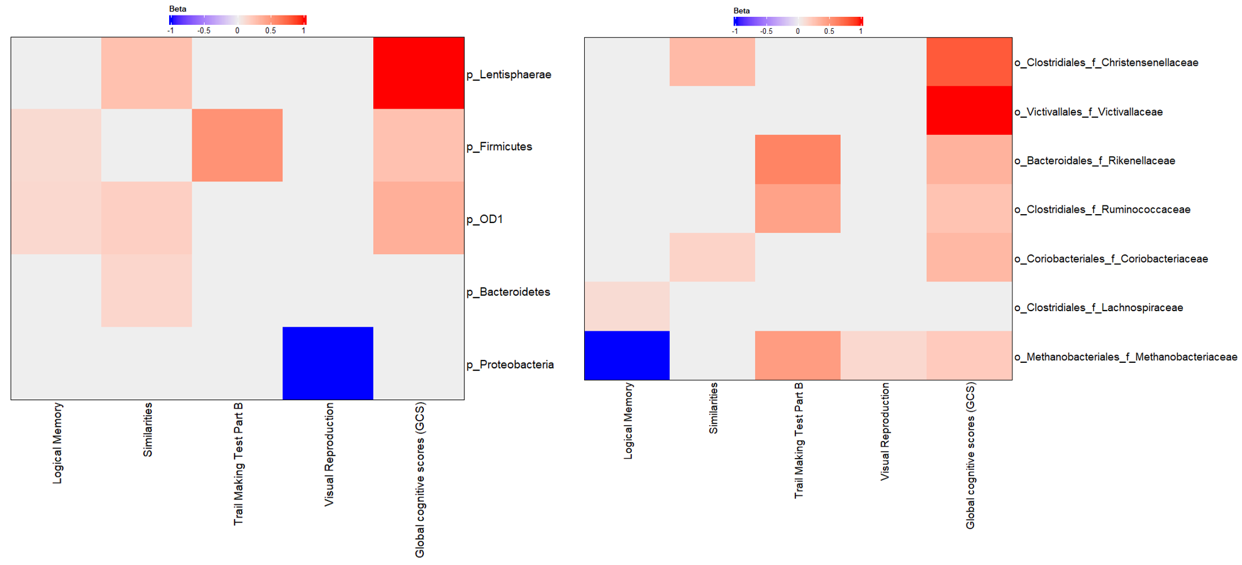

**Figure S7.** Multivariable association between gut microbiota and global cognitive scores. Heatmap depicting the significant (adjusted p-value < 0.05) taxa associated with GCS along with scores of tests composing GCS at the phylum (left) and family (right).

**Gut microbiome mediated the association of LE8 with global cognitive scores**

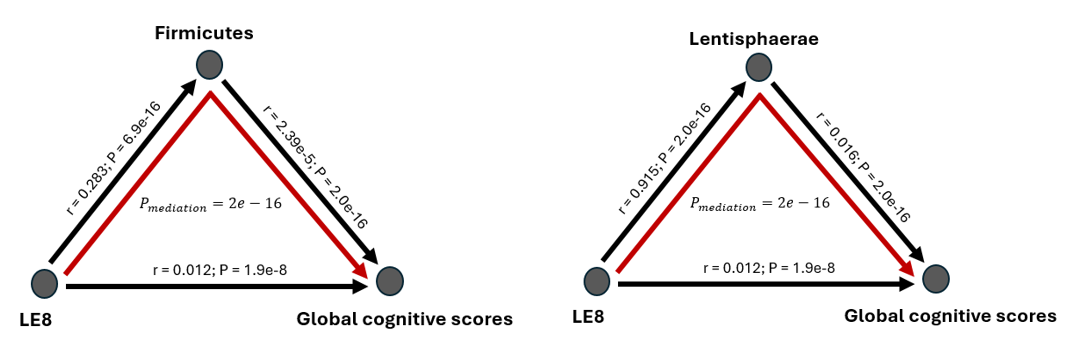

**Figure S8.** Mediation analysis of gut microbiome on the association between LE8 adherence scores and global cognitive scores. The mediation effect of bacteria *Firmicutes* and *Lentisphaerae* on the association between moderate LE8 adherence with lower cognition.

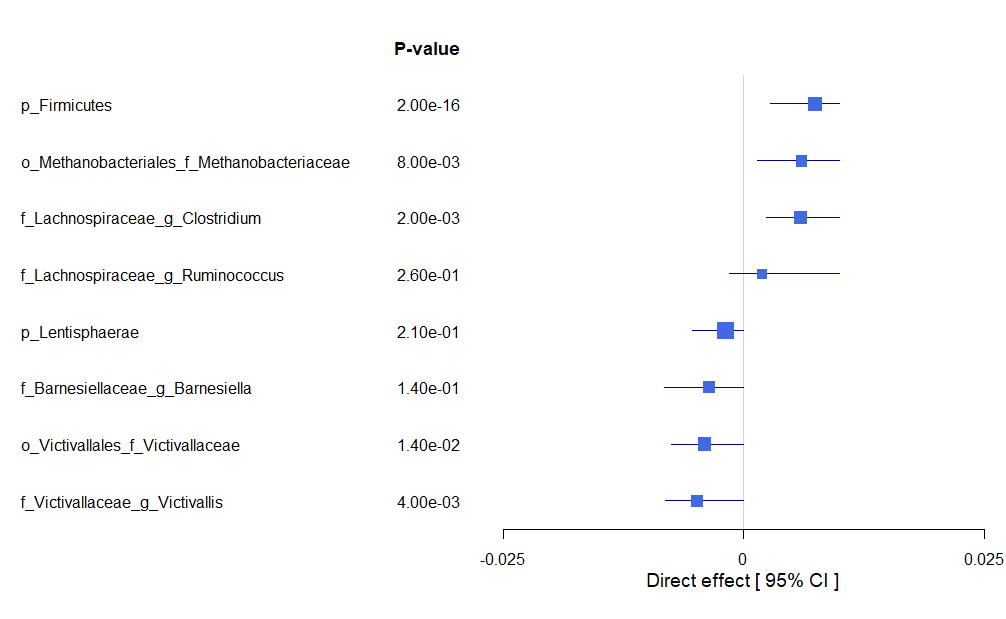

**Figure S9.** Forest plot showing the estimated direct effects.
